# mTOR drives cerebrovascular dysfunction and blood-brain barrier breakdown in a model of Alzheimer’s disease with cerebral amyloid angiopathy

**DOI:** 10.64898/2026.06.23.733858

**Authors:** Candice E. Van Skike, Lauren R. Miller, Stephen F. Hernandez, Stacy A. Hussong, Haneen Makhlouf, Aishwarya Chitoor Muppala, Nicholas DeRosa, Jordan B. Jahrling, Kelly T. Dineley, Veronica Galvan

## Abstract

Cerebral amyloid angiopathy (CAA) is characterized by the deposition of amyloid β fibrils (Aβ) within walls of the cerebrovasculature and contributes to intracerebral hemorrhage, ischemic stroke, and cognitive dysfunction in patients with Alzheimer’s disease (AD) and in non-pathological aging. Previous studies have shown that mTOR drives cerebrovascular dysfunction and cognitive impairment observed in AD, vascular cognitive impairment, and normative aging. However, the mechanisms by which mTOR contributes to CAA are unknown. Here, we show that mTOR drives the accumulation of fibrillar vascular Aβ lesions in the Tg2576 Model of AD with CAA (using equal numbers of female and male mice), which directly impair endothelium-dependent cerebrovascular reactivity. Additionally, we found that blood-brain barrier (BBB) breakdown and remodeling of tight junction proteins, dependent on mTOR, are associated with increased cerebral microhemorrhages. Finally, we show that mTOR contributes to neurovascular uncoupling in Tg2576 AD mice through nNOS dysfunction and inhibition of non-nitric oxide synthase-dependent contributions to neurovascular coupling (NVC). Contextual memory impairments were ameliorated by the mTOR inhibitor rapamycin. Improvements in memory were associated with reduced cerebrovascular Aβ fibril accumulation, enhanced endothelium-dependent vasodilation, reduced fibrillar Aβ load, restoration of BBB integrity, attenuation of intracerebral microhemorrhage, and restoration of NVC. These data indicate that mTOR drives vascular accumulation of fibrillar Aβ, including those associated with brain vasculature, and mediates cerebrovascular dysfunction in a model of AD with CAA. Thus, mTOR inhibitors represent a promising treatment option for patients with CAA and AD.

## Introduction

The Alzheimer’s Disease Neuroimaging Initiative (ADNI) determined that cerebrovascular deficits arise as the earliest and most profound injury in Alzheimer’s disease (AD) progression (Iturria-Medina et al., 2016), preceding contextual memory impairment, brain atrophy, deposition of pathogenic amyloid β (Aβ) and tau, and clinical diagnosis of AD (Iturria-Medina et al., 2016, Hays et al., 2016, Luo et al., 2023). Cerebral amyloid angiopathy (CAA), defined as the deposition of Aβ within the walls of brain blood vessels, is a major cerebrovascular pathology that occurs in approximately 90% of AD patients and is also found in elderly adults without AD albeit at lower rates (Yamada et al., 1987, Fisher et al., 2022). Aβ However, the mechanisms that contribute to the development of CAA are not well understood.

Similar to hAPP(J20) (Hsia et al., 1999, Mucke et al., 2000, Spilman et al., 2010, Lin et al., 2013) and 3xTg-AD mice (Oddo et al., 2003), Tg2576 mice (Hsiao et al., 1996, Taglialatela et al., 2009, Comery et al., 2005) recapitulate the cognitive and synaptic deficits of Alzheimer’s Disease (AD), but also develop extensive vascular fibrillar Aβ deposits that mimic the vascular lesions found in cerebral amyloid angiopathy (CAA) patients(Yamada et al., 1987). CAA is associated with a significantly higher risk of dementia and intracranial hemorrhage. Cerebrovascular fibrillar Aβ deposition begins at ∼9-12 months of age in Tg2576 mice (Hsiao et al., 1996, Kawarabayashi et al., 2001, Domnitz et al., 2005) and is correlated with vascular dysfunction (Han et al., 2015, Tarantini et al., 2017a) and increased occurrence of microhemorrhages (Yakushiji et al., 2020, Hur et al., 2018, Han et al., 2015). These phenotypes make Tg2576 mice an ideal model for studying CAA-related vascular dysfunction in AD.

The mammalian/mechanistic target of rapamycin (mTOR) has been identified as a critical mediator of cerebrovascular dysfunction (Van Skike et al., 2018, Van Skike et al., 2020, Van Skike et al., 2021, Lin et al., 2017, Lin et al., 2013, Jahrling et al., 2018) in models of AD (Lin et al., 2013, Lin et al., 2017, Van Skike et al., 2018), vascular cognitive impairment (Jahrling et al., 2018, Van Skike et al., 2018), and in normative aging (Van Skike et al., 2020). These studies have shown that attenuation of mTOR activity with rapamycin prevents AD-and age-associated cerebrovascular dysfunction through the restoration of the endothelial and neuronal forms of nitric oxide synthase (eNOS and nNOS, respectively)(Van Skike et al., 2021). To further establish the mTOR-dependent regulation of cerebrovascular dysfunction, we assessed vascular reactivity, blood-brain barrier (BBB) integrity, neurovascular coupling, and cognitive function in a model of cerebral amyloid angiopathy, which is characterized by extensive fibrillar Aβ lesions within the brain vasculature. We found that mTOR promotes accumulation of vascular fibrillar Aβ lesions in Tg2576 mice. *In vivo* two photon imaging indicated that fibrillar Aβ lesions were associated with impaired local vascular reactivity, and mTOR attenuation with rapamycin preserved local vascular reactivity even in vascular segments with high Aβ load. Given that CAA is associated with an increased incidence of intracranial hemorrhage, Tg2576 mice exhibited BBB breakdown as measured using *in vivo* two photon microscopy as well as microhemorrhages detected in tissue sections. BBB integrity was restored with the mTOR inhibitor rapamycin and resulted in reduced incidence of microhemorrhages through changes in tight junction protein expression. Neurovascular coupling (NVC) in response to whisker pad stimulation was significantly impaired in Tg2576 mice. NVC was restored to wildtype levels by mTOR attenuation, suggesting that mTOR is a mechanistic driver of the NVC functional impairment. Finally, we found that Tg2576 mice had significantly impaired contextual memory that was ameliorated to levels indistinguishable from wildtype with the inhibition of mTOR. Together, our data support the role of mTOR in the development of CAA, cerebrovascular dysfunction, and cognitive impairment in a model of AD with CAA.

## Materials and Methods

### Animals and experimental diet

We performed experiments using 2 separate cohorts of male and female wild-type (WT) littermate mice and heterozygous Tg2576 transgenic mice (RRID:IMSR_TAC:1349) that overexpress mutant APP (isoform 695) with the Swedish (KM670/671NL) mutation under the control of the hamster prion promoter, resulting in elevated levels of Aβ, parenchymal amyloid plaques, and vascular amyloid angiopathy associated with cognitive impairment. Mice were bred at the University of Texas Medical Branch (UTMB, Galveston, TX, USA) animal care facility by mating hemizygous Tg2576 males with B6SJL/F1J females, as previously described (Hsu et al., 2017, Jahrling et al., 2014). Transgenic and non-transgenic female and male mice (WT, control) were randomly assigned to receive chow supplemented with either microencapsulated rapamycin (14 ppm, Emtora Biosciences, San Antonio, TX, USA) or vehicle-supplemented chow (eudragit, Emtora Biosciences) as previously described (Pierce et al., 2013, Van Skike et al., 2018, Spilman et al., 2010, Harrison et al., 2009, Halloran et al., 2012, Van Skike et al., 2021, Van Skike et al., 2023). This diet formulation provided the animals with an estimated 1.65 mg/kg/day of rapamycin, which was calculated using the average amount of food consumed per mouse within a group-housed cage (≤ 5 animals/cage) during these studies. Mice were treated with experimental diets starting at 4 months of age to the end of study. Animals had continuous access to diets and water. All studies were approved by the University of Texas Health Science Center San Antonio’s (UTHSCSA) Institutional Animal Care and Use Committee (Animal Welfare Assurance Number A3345-01) and complied with ARRIVE guidelines. Equal numbers of female and male mice were used in each group (5-6 mice) in both Cohort 1 and 2 (**Figure 1**). Minimum group size was calculated based on pilot two-way ANOVA data from the most variable measure, baseline neurovascular coupling responses (described in later methods section), n = 4 per group provides 95% power to detect a genotype main effect corresponding to 50% explained variance (partial η² = 0.5; Cohen’s f = 1.0) at α = 0.05 in two-way ANOVA. Outcomes were measured in a blinded manner

**Figure 1.**
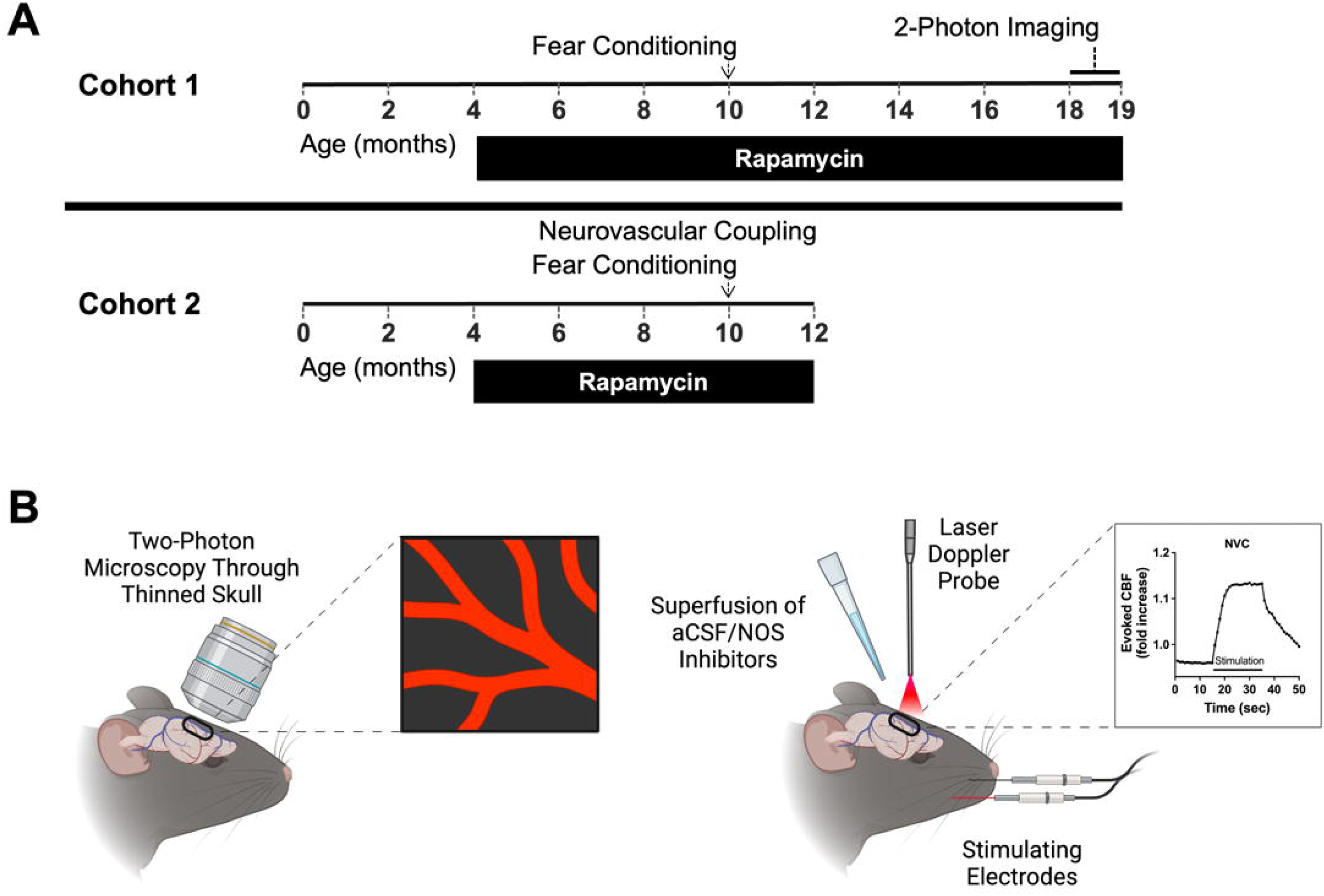
Experimental overview of the two Tg2576 cohorts, contextual fear conditioning, two-photon imaging, and neurovascular coupling. **(A)** Timeline of events for contextual fear conditioning, neurovascular coupling (NVC), and two-photon imaging in the Tg2576 cohorts. In both cohorts, rapamycin or vehicle treatment began at 4 months of age. Hippocampal-dependent contextual fear conditioning was conducted at 10 months of age in both cohorts and data were combined for the final analysis. In cohort 1, *in vivo* two-photon imaging was performed between 18-19 months of age. Neurovascular coupling (NVC) was performed in the second cohort of Tg2576 mice at 12 months of age. **(B)** Graphical representation of *in vivo* two-photon microscopy of the barrel cortex through intact, thinned bone. **(C)** Graphical representation of NVC using laser Doppler flowmetry to measure cerebral blood flow (CBF) in response to electrical whisker pad stimulation. Panels B and C were created in BioRender. Makhlouf, h. (2025) https://BioRender.com/oso6dlx.

### In vivo two-photon imaging

#### Surgical thinning of mouse skulls

18-month-old transgenic and wildtype Tg2576 mice were anesthetized and maintained under 2.0-2.4% isoflurane gas (Butler Schein, Dublin, OH, USA). The mice were placed on a custom fabricated water-warmed heating platform to maintain core body temperature. Hair was removed from the top of the mouse’s skull and the exposed skin was swabbed with iodine and isopropanol. A small 1.5 cm sagittal incision was made over the midline of the skull, the skin gently retracted, and a custom fabricated metal stereotaxic holder was secured to the dry skull with Loctite professional liquid super glue. The holder was then fastened to a stereotaxic ear bar holder mounted to a custom heating platform, both adapted for use with the 2-photon microscope assembly. The skull over the somatosensory cortex (1-3mm AP from bregma, 2-4mm lateral) was thinned by a variable speed electric drill with a small burr bit (Fine Science Tools, Foster City, CA, USA), followed by further thinning and smoothing with a surgical razor blade to achieve a thickness of about 50 microns. The possibility of heat injury was reduced by frequent application of sterile saline and constant moving of the drill bit during the thinning process. A 1.5mm round glass coverslip was glued over the thinned skull using Loctite brush-on super glue. When dry, the anesthetized mouse was transported to the microscopy core for imaging.

#### Two-photon imaging of Aβ plaques, BBB integrity, and cortical vasodilation

Transcranial imaging was performed using the Prairie Two-Photon System equipped with a Nikon Eclipse SN1 upright microscope and acoustic optical deflection mode. A 25X APO water immersion objective with NA 1.10 was used to collect all images. The Ti:Sapphire laser was tuned at each use to 800 nm. Mice were injected with 2.0 mg/kg methoxy-X04 (4920, Tocris, Avonmouth, England, UK), a blood-brain barrier permeable fibrillar Aβ fluorescent probe, dissolved in sterile saline and 3% Kolliphor (C5135, Sigma, St. Louis, MO, USA) the day before they were imaged. After several imaging planes were acquired for methoxy-labeled Aβ, the vasculature was illuminated by administering 100 μL of 10 mg/mL Texas Red 70kD dextran (D1830, Invitrogen, Waltham, MA, USA) in sterile saline via the tail vein and baseline images were acquired. Thirty minutes later, another set of images was acquired to measure BBB breakdown as extravasation of the fluorescent dextran. Afterward, acetylcholine (ACh, 10μg/kg, i.p., A6625, Sigma) was administered to elicit endothelium-dependent vasodilation and images were collected 10 minutes after the injection.

### Two photon image analysis and statistics

#### Vascular Aβ burden

To assess the extent of vascular amyloid burden, images were analyzed in ImageJ (NIH, Bethesda, MD, USA). The z-stack range was adjusted to span the diameter of the vessel of interest for each mouse and compiled with a maximum intensity projection. An index of vessel associated Aβ was calculated by measuring the area of methoxy-X04 labeling from the green channel and dividing by the area of the vasculature seen in the red channel. These numbers were normalized to the wildtype group to control for background methoxy-X04 staining. Since wildtype animals do not have significant Aβ deposits, they were excluded from analysis. A t-test was performed between rapamycin-and vehicle-treated Tg2576 animals to compare the levels of vascular-associated Aβ.

#### Brain Aβ fluorescence

We assessed the fluorescence of both vessel-associated and parenchymal Aβ deposits to test the hypothesis that TOR drives vascular Aβ aggregation. To do this, the fluorescence of multiple continuous methoxy-X04 stained (green), Aβ segments were measured with ImageJ, and corrected using the formula reported in McCloy et al. (2014) and then normalized to WT. This formula calculates the corrected integrated density as “integrated density – (area of interest × background fluorescence).” Five to eight measurements were made for each image, depending on the number of continuous vascular Aβ segments present in the image. Whenever possible, continuous vascular Aβ segments were measured as one unit so the analysis would not be skewed by oversampling from similar areas, and the mean fluorescent intensity of the continuous area was used in the equation given above. Separate t-tests were performed between rapamycin-and vehicle-treated transgenic Tg2576 animals to compare vascular and parenchymal Aβ fluorescence.

#### BBB integrity measured as 70kD dextran extravasation

Z-stack images spanning 50 μM were acquired of the Texas Red 70kD dextran and assembled in Image J. Mean background fluorescence was measured at 5 minutes and 30 minutes, and the percent increase in background fluorescence relative to the 5 min baseline, indicative of extravasation, was calculated for each mouse.

#### Endothelial-dependent vasodilation

Vessel diameter (stained with Texas Red dextran) was measured with ImageJ by manually drawing straight lines across the vessel of interest in the images acquired 30 minutes after dextran and 10 minutes after acetylcholine injection (10μg/kg, i.p., A6625, Sigma). Vasodilation is expressed as percent change from the pre-acetylcholine baseline diameter. 10 measurements were made per animal and the presence or absence of Aβ was noted. A 2-way repeated measures ANOVA was performed to assess vasodilatory capacity based on treatment and the presence of fibrillar Aβ plaques. Sidak’s multiple comparisons test was performed to compare the effect of Aβ presence between the treatment groups. Since wildtype animals do not have significant vascular amyloid burden, the analysis of the association of Aβ burden and vasodilation was only performed in Tg2576 animals.

### Microhemorrhage quantification in tissue sections

10μm coronal cryosections from snap-frozen hemibrains were stained with H&E by the San Antonio Nathan Shock Center Pathology Core to localize and quantify acute cerebral microhemorrhages (Liu et al., 2014, Sumbria et al., 2018). Cerebral microhemorrhages were counted at 10X magnification by an observer blinded to experimental condition. Two-way ANOVAs (Genotype x Treatment) were performed to detect group differences in the number of microbleeds.

### Immunofluorescent quantification of tight junction proteins and MMP9

Quantification of tight junction proteins via immunofluorescent detection in frozen tissue was performed as previously described (Van Skike et al., 2021). Briefly, cortical cryosections were cut from hemibrains at ten microns and fixed in 4% PFA for 30 minutes. Sections were washed with TBS and subsequently incubated in blocking solution (5% BSA and 5% goat serum in TBS) for 1 hour at room temperature. Tissue sections were incubated with primary antibodies against claudin-5, ZO-1, and MMP-9 applied in conjunction with DyLight tomato lectin incubated overnight at 4°C (see **Table 1** for antibody information). After washing, tissue sections were incubated with Alexa Fluor 594-conjugated goat anti-rabbit secondary for 1 hour, counterstained with DAPI (D1306, Thermo Fisher Scientific, Waltham, MA, USA), and slides were mounted with ProLong Gold Antifade Mountant (P36930, Thermo Fisher Scientific, Waltham, MA, USA).

**Table 1:** Antibodies.

| Antibody Target | Catalog #/Vendor | Concentration | RRID | References |
| --- | --- | --- | --- | --- |
| Alexa 488-lectin | DL1174, Vector Laboratories | 1:750 | RRID: AB_2336404 | (Van Skike et al., 2018) |
| Alexa 594-anti-rabbit | A-11012, Thermo Fisher Scientific | 1:500 | RRID: AB_2534079 |  |
| Claudin-5 | Catalog no. 352500, Life Technologies | 1:500 | RRID: AB_2533200 | (Van Skike et al., 2018) |
| MMP-9 | PA5-13199, Invitrogen | 1:50 | RRID: AB_2144757 |  |
| ZO-1 | 61-7300, Invitrogen | 1:50 | RRID: AB_2533938 | (Zhang et al., 2022) |

Images were collected using a Zeiss LSM 780 NLO confocal microscope equipped with a 40X water immersion objective and numerical aperture of 1.1. Positive fluorescent immunoreactivity was quantified in the original grayscale images by a researcher blinded to treatment condition as mean fluorescence intensity using Image J Fiji (Schindelin et al., 2012). After analysis, images were colorized and processed in Adobe Photoshop while maintaining identical contrast and brightness ratios. Tissue from WT controls were included in all experiments to assess the specificity of immunoreactivity. Data were analyzed using a two-way ANOVA, followed by Tukey’s post hoc tests.

### Neurovascular coupling

Neurovascular coupling (NVC) was measured in 12-month-old mice using laser Doppler flowmetry based on previously reported methods (Royea et al., 2017, Van Skike et al., 2021), by an experimenter blinded to genotype and treatment condition. Animals were anesthetized with a combination of ketamine and xylazine (100 mg/kg and 12.5 mg/kg, respectively) and placed in a stereotaxic frame on a heating pad to maintain body temperature at 37°C. Blood oxygen saturation, heart rate, respiration rate, and blood pressure were continuously monitored using the PhysioSuite (Kent Scientific, Torrington, CT, USA). Measurements of baseline NVC and NVC with nNOS inhibited were performed in less than 20 minutes, a time during which mouse physiological parameters, including blood gases, are stable (Tong et al., 2012, Ongali et al., 2014). The bone over the left somatosensory barrel cortex (0.5-1.5 mm posterior and 3-4.5 mm lateral from Bregma) was removed, a silicone elastomer barrier (World Precision Instruments, Sarasota, FL, USA) was made around the craniotomy, artificial CSF (aCSF, 3525, Tocris) was continuously superfused over the craniotomy, and a laser Doppler probe (Transonic, Ithaca, NY, USA) was positioned over the barrel cortex. NVC was measured as increased cerebral blood flow (CBF) relative to baseline in response to contralateral electrical whisker pad stimulation (20 seconds at 5 Hz and 1mA, with 40 second intervals). CBF was measured using LabChart (ADInstruments, Colorado Springs CO, USA) and changes in CBF are expressed as fold increase relative to average baseline CBF collected during the 15 seconds pre-and post-stimulus.

We have previously utilized *N*^ω^-Propyl-L-arginine (L-NPA) and *N*^ω^-Nitro-L-arginine methyl ester hydrochloride (L-NAME) to define the contributions of nNOS and other NOS (namely eNOS) to NVC (Van Skike et al., 2021). Prior studies have shown that L-NPA has 150-fold selectivity for the inhibition of nNOS over eNOS (Zhang et al., 1997) and has been utilized as a specific nNOS inhibitor *in vivo* (Beamer et al., 2012, Sousa et al., 2015, Bonvento et al., 2000, Burke and Buhrle, 2006, Stefanovic et al., 2007, Yang et al., 1999, Almeida et al., 2018, Kato et al., 2009, Lubomirov et al., 2017). although there are reports that L-NPA lacks nNOS specificity in models that utilize *ex vivo* tissue with concentrations of L-NPA up to 1000X higher than the Ki for nNOS (Overend and Martin, 2007, Pigott et al., 2013). To verify the specificity of L-NPA, we have previously demonstrated that superfusion of 200nM L-NPA does not have any impact on endothelium-dependent, ACh-induced vasodilation, which is mediated by eNOS, in WT mice (Van Skike et al., 2021), thus ruling out non-specific actions of 200 nM L-NPA on eNOS-dependent activity. Further, we have demonstrated that the addition of the general NOS inhibitor L-NAME at 10 μM produces a near-complete inhibition of endothelium-dependent vasodilation induced by subsequent superfusion of ACh (Van Skike et al., 2021).

After measuring baseline NVC in the presence of aCSF, to effectively and specifically inhibit nNOS, we utilized a 10 minute continuous superfusion of 200 nM L-NPA (80587, Cayman Chemical, Ann Arbor, MI, USA) prior to additional NVC measurements (Zhang et al., 1997). To define the contribution of eNOS to NVC, after superfusion with 200 nM L-NPA, we continuously superfused a combination of 200 nM L-NPA and 10 μM L-NAME (80210, Cayman Chemical), a non-selective NOS inhibitor, for 10 minutes prior to additional NVC measurements. Since nNOS activity had been previously blocked with L-NPA, additional decreases in NVC observed after the addition of L-NPA + L-NAME defined the specific contribution of eNOS to NVC. Potential contributions of the inducible form of NOS (iNOS) to NVC are unlikely since it is constitutively activated once assembled (Stuehr, 1999).

The average of 5 stimulations for each of the 3 conditions examined (baseline (aCSF, vehicle), L-NPA, and L-NPA + L-NAME) were plotted for each animal and the area under the curve during stimulation, with baseline normalized to one, was calculated. The nNOS-dependent, the non-nNOS, L-NAME-sensitive (eNOS)-dependent, and the NOS-independent contributions to NVC were calculated as (1) the percent reduction in NVC by L-NPA compared to baseline vehicle stimulation, (2) the percent reduction in NVC by subsequent L-NPA + L-NAME application relative to vehicle stimulation, subtracting the percent reduction by L-NPA alone to isolate the unique contribution of eNOS, and as (3) the remaining magnitude of the NOS-independent NVC response after subtraction of the percent reduction by L-NPA alone and by L-NPA + L-NAME respectively. Data were analyzed with a two-way ANOVA followed by Tukey’s post hoc test.

### Contextual Fear Conditioning

To measure hippocampal-dependent contextual memory, all animals underwent contextual fear conditioning at 10 months of age. Briefly, mice were acclimated to the testing environment for 4 hours prior to being placed into test cages within isolation chambers (Coulbourn Instruments, Holliston, MA, USA) and exposed to two pairings of a 30s auditory stimulus (white noise), which co-terminated with a 2 second foot shock (1mA). 24-hour recall of hippocampal-dependent contextual memory was assessed during a 5-minute exposure to the context itself, conducted in the absence of any tone or shock. FreezeFrame 4 software (Actimetrics, Lafayette, IN, USA) was used to record, monitor, and quantify freezing behavior. Percent time spent freezing was calculated for each mouse and mice were excluded if less than 5% freezing, indicative of hyperactivity within this strain (Gil-Bea et al., 2007, Lalonde et al., 2012), was detected during training. This resulted in the exclusion of 7 animals from this analysis: 6 from the Tg2576+Vehicle group and 1 from the Tg2576+Rapamycin group. One additional animal was excluded for the WT+Vehicle group due to complete lack of freezing observed during the contextual test. Data were analyzed using a two-way ANOVA followed by Tukey’s multiple comparison test among all means.

## Results

### Attenuation of mTOR selectively reduces vascular amyloid β burden

Previous studies have demonstrated the reduction of parenchymal Aβ deposition via pharmacological inhibition of mTOR with chronic rapamycin treatment in hAPP(J20) mice (Lin et al., 2013, Spilman et al., 2010) and 3xTg-AD mice (Majumder et al., 2011) or genetic reduction of mTOR in Tg2576 mice (Caccamo et al., 2014). Furthermore, inhibition of mTOR by rapamycin also decreased the vascular Aβ burden in hAPP(J20) mice (Lin et al., 2013). To determine whether mTOR may regulate accumulation of fibrillar Aβ in the cerebrovasculature of Tg2576 mice, we used methoxy-X04 labeling and *in vivo* transcranial two-photon imaging to measure cerebrovascular and parenchymal Aβ load in 18-month-old Tg2576 mice. Extensive vascular fibrillar Aβ accumulation (**Figure 2A**) was observed in vehicle-treated Tg2576 animals. Congruent with previous studies that showed reduction of parenchymal Aβ load by mTOR attenuation (Caccamo et al., 2014, Lin et al., 2013, Majumder et al., 2011, Spilman et al., 2010), mTOR attenuation with rapamycin reduced the total area of vascular fibrillar Aβ lesions in 18-month-old Tg2576 mice by 40.6% compared to vehicle-treated controls (**Figure 2A, B**). Additionally, rapamycin treatment reduced the amount of vessel-associated fibrillar Aβ by 28.9% (**Figure 2A, C**). This reduction was specific to vascular Aβ lesions, as the density of parenchymal fibrillar Aβ lesions was not changed with mTOR inhibition in Tg2576 mice (**Figure 2D, E, F**).

**Figure 2.**
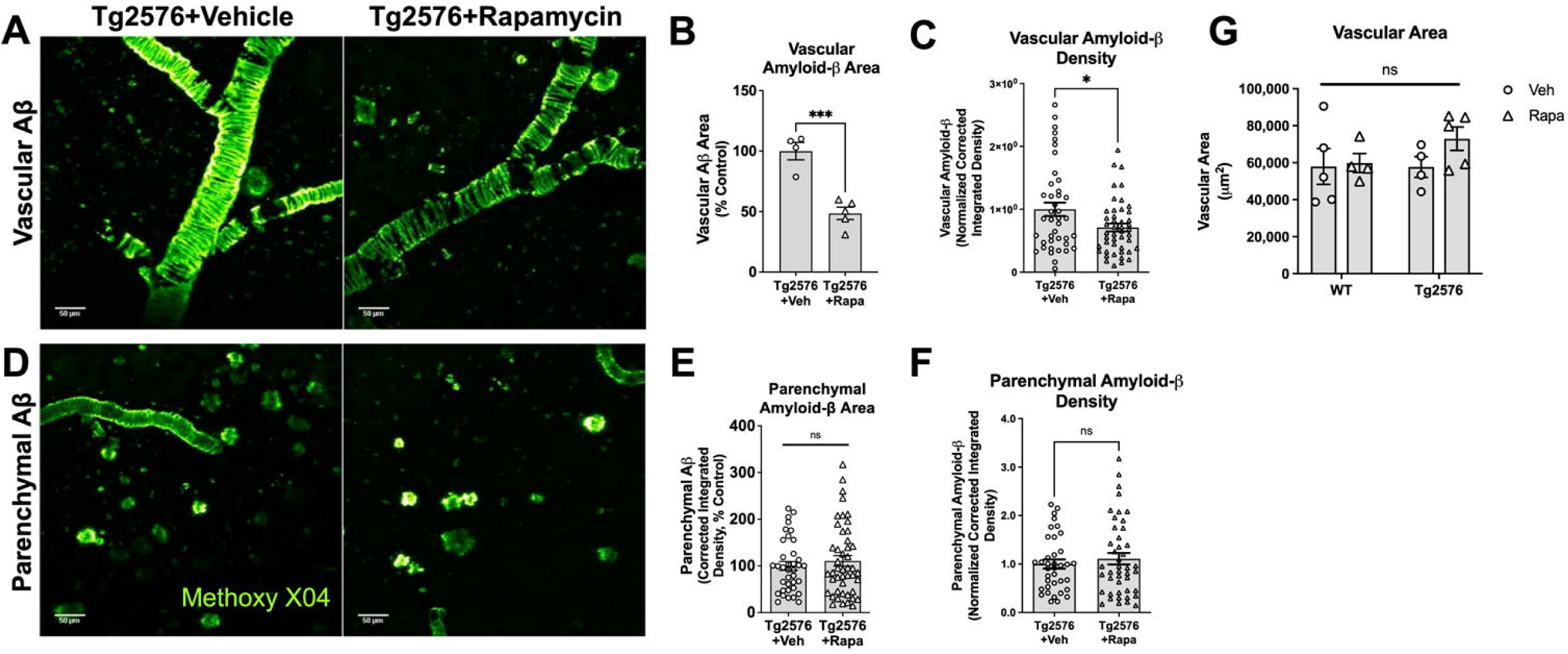
**mTOR contributes to vascular amyloid-**β **accumulation in the Tg2576 model of AD with cerebral amyloid angiopathy. (A)** Representative z-stack projections generated with *in vivo* two photon microscopy showing the extent of fibrillar vascular Aβ lesions in 18-month-old transgenic Tg2576 mice treated with rapamycin or vehicle. Fibrillar Aβ is visualized with Methoxy X-04 (green). **(B)** Chronic mTOR inhibition significantly reduces vascular Aβ burden as measured by Aβ-positive area relative to total area of the vasculature in 18-month-old Tg2576 mice (*** indicates t(7)=5.950, p=0.0006). Data represent mean±SEM of n=3-5 per group. **(C)** Density of vascular fibrillar Aβ lesions in Tg2576 mice is reduced by chronic rapamycin to attenuate mTOR (*, t(85)=2.39, p=0.019). Data represent mean±SEM of n=40-46 total measurements of continuous CAA segments from 3-5 unique fields of view from 3-5 mice per group. **(D)** Representative in vivo two-photon microscopy images of parenchymal Aβ lesions in transgenic Tg2576 mice treated with vehicle or rapamycin. **(E)** Attenuation of mTOR does not affect parenchymal Aβ accumulation in the Tg2576 mouse model of Alzheimer’s disease (t(82)=0.19, p=0.85). Data represent mean±SEM of n=37-46 total measurements of individual parenchymal Aβ plaques from 3-5 unique fields of view from 3-5 mice per group. **(F)** Total brain vascular area within the imaging region is unchanged in Tg2576 mice relative to non-transgenic (WT) littermates (F(1,14)=0.76, p=0.40), and is unchanged with chronic mTOR attenuation via rapamycin (F(1,14)=1.34, p=0.26). Data represent mean±SEM of n=3-5 per group. Representative images of brain vasculature shown in Figure 2A.

In contrast to previous findings of preserved vascular density by rapamycin in younger hAPP(J20) mice modeling AD (Lin et al., 2013), we found vascular density within the imaging window was not altered by genotype nor rapamycin treatment (**Figure 2G**), indicating that the reduction in vascular Aβ accumulation (**Figure 2A, B**) was due to a specific decrease of vascular Aβ accumulation by mTOR attenuation without being driven by a change in vascular density. Taken together, these data indicate that mTOR specifically drives accumulation of cerebrovascular fibrillar Aβ lesions in the Tg2576 mouse model of AD.

### Chronic mTOR attenuation preserves cerebrovascular reactivity in vascular segments with high Aβ load

Prior studies have shown impaired vascular reactivity in hAPP(J20) (Tong et al., 2009, Nicolakakis et al., 2008, Ongali et al., 2014, Royea et al., 2017) and Tg2576 mice (Han et al., 2015). To determine a potential role of mTOR in driving impaired vasomotor responses associated with CAA-like cerebrovascular fibrillar Aβ deposition in Tg2576 mice (Han et al., 2015), we measured vasodilatory responses to acetylcholine (ACh) in contiguous segments of vasculature with (Aβ^+^) and without (Aβ^-^) fibrillar Aβ deposits. Chronic mTOR attenuation with rapamycin significantly enhanced vasodilatory responses to ACh in cortical vasculature of WT mice (p=0.0013, **Figure 3A, B)**. Furthermore, mTOR attenuation significantly improved vasodilatory responses in Aβ^-^and in Aβ^+^ vessel segments of transgenic Tg2576 mice compared to Tg2576 vehicle-treated in Aβ^+^ vessel segments (*** indicates p=0.0002, * indicates p=0.014; Figure 3A, B). Severe impairment of vasodilatory responses to ACh in Aβ^+^ vessel segments from vehicle-treated Tg2576 mice (**Figure 3A, B**), however, was negated by mTOR attenuation, such that the magnitude of dilatation in Aβ^+^ vessel segments in rapamycin-treated Tg2576 mice was nearly indistinguishable from that of Aβ^-^vessel segments in vehicle-treated Tg2576 mice (**Figure 3B**). Thus, mTOR attenuation profoundly rescues vascular reactivity impairments associated with CAA-like lesions in cerebrovasculature of Tg2576 mice.

**Figure 3.**
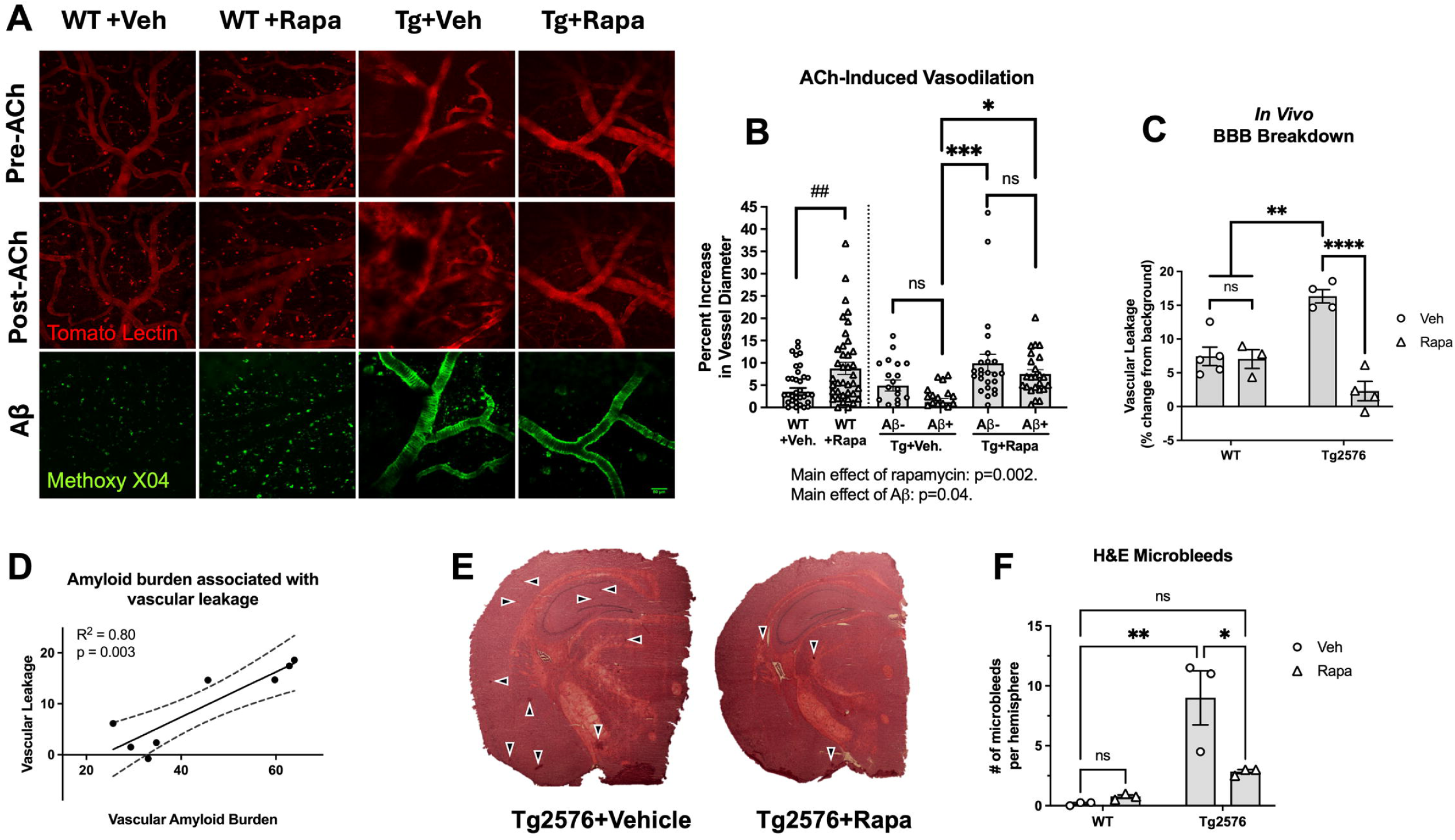
Vascular reactivity and BBB integrity is restored by chronic mTOR inhibition in the presence of high fibrillar. **A**β **load. (A)** Representative *in vivo* two-photon images showing dextran extravasation indicating BBB breakdown and endothelium-dependent vasodilation (middle row) as a function of original vessel diameter (top row). Vasculature was illuminated with i.v. Texas Red 70kD dextran and measurements were taken from vessel segments with and without fibrillar Aβ (Methoxy-X04, green, bottom row). **(B)** Endothelium-dependent vasodilation was elicited by acetylcholine (ACh) and vascular reactivity was measured as % increase in vessel diameter relative to baseline diameter. Chronic mTOR attenuation improved vascular reactivity in WT mice (two-tailed unpaired t test, t(79)=3.332, p=0.0013). Chronic mTOR inhibition increases vascular reactivity (main effect of rapamycin: q(85)=15.51, p=0.0002, two-way ANOVA). Additionally, vascular reactivity is impaired by the presence of fibrillar Aβ (main effect of Aβ: q(85)=4.57, p=0.04, two-way ANOVA) in transgenic Tg2576 mice. Preservation of vascular reactivity by rapamycin is especially apparent in vessel segments with fibrillar Aβ deposits (* indicates Tukey’s posthoc comparing Aβ+ vessel segments in Tg2576+Vehicle vs Tg2576+Rapa: q(85)=4.4, p=0.014), indicating that attenuation of mTOR preserves vascular reactivity even in the presence of high Aβ load. There is also a significant increase in vascular reactivity Aβ-vessel segments of Tg2576+Rapamycin versus Aβ+ vessel segments in Tg2576+Vehicle mice (*** indicates Tukey’s posthoc comparison, q(85)= 6.10, p=0.0002). Data represent mean±SEM of 5 unique Aβ+ and 5 unique Aβ-vessel segments of n=3-5 per condition. **(C)** Blood-brain barrier breakdown in 18-month-old Tg2576 mice is significantly reduced with chronic mTOR inhibition (** significant increase over both WT treatment groups, Two-way ANOVA with Tukey’s posthoc, q(12)<7.02, p<0.003; **** q(12)=10.52, p<0.0001). **(D)** There is strong direct correlation between vascular Aβ burden (Figure 2) and BBB breakdown in Tg2576 mice (r=0.80, p=0.003). Data are overlaid on a solid regression line of best fit with dotted lines representing the standard error. **(E)** Representative images of H&E staining for microbleeds (indicated by black arrows). **(F)** The number of microbleeds per hemisphere is significantly increased in Tg2576 mice (** Tukey’s q(8)=7.79, p=0.003 versus WT+Vehicle). The number of microbleeds is significantly reduced in Tg2576 mice treated with rapamycin (* q(8)=5.44, p=0.02 versus Tg2576+Vehicle) to a level that is not significantly different from control (q(8)=2.35, p=0.40, n.s.). Data represent mean ±SEM of n=3 mice per group.

### Attenuation of mTOR preserves BBB integrity and reduces the incidence of microhemorrhages

Prior studies have demonstrated mTOR drives BBB breakdown in the hAPP(J20) model of AD (Lin et al., 2013, Lin et al., 2017, Van Skike et al., 2018) and in models of vascular cognitive impairment (Van Skike et al., 2018, Jahrling et al., 2018). To determine if mTOR regulates BBB integrity in the Tg2576 model of AD with extensive CAA-like amyloid deposits, we utilized transcranial *in vivo* two-photon microscopy to measure extravasation of intravenously administered fluorescently-labeled 70kD dextran in 18-month-old animals (Figure 3A, C). Similar to previous reports utilizing tissue sections from Tg2576 mice (Biron et al., 2011, Ujiie et al., 2003), we identified significant BBB breakdown in vehicle-treated Tg2576 mice using *in vivo* two-photon microscopy (**Figure 3A, C**). Consistent with a role of mTOR in BBB breakdown (Van Skike et al., 2018), chronic attenuation of mTOR with rapamycin significantly reduced dextran extravasation, showing preserved of BBB integrity in Tg2576 mice similar to WT controls (**Figure 3C**). We examined the relationship between BBB breakdown and vascular fibrillar Aβ burden and identified a strong positive correlation between BBB breakdown and vascular Aβ load in Tg2576 mice (r=0.80, p=0.003, **Figure 3D**), suggesting that fibrillar vascular Aβ lesions are associated with BBB dysfunction.

CAA is associated with spontaneous, recurring intracerebral microbleeds in older adults (Pezzini and Padovani, 2008, Mehndiratta et al., 2012) and in AD (Chin et al., 2024, Akoudad et al., 2016, Benedictus et al., 2013). These microbleeds are likely related to BBB breakdown (Freeze et al., 2018, Freeze et al., 2019, van Nieuwenhuizen et al., 2017, Hartz et al., 2012), thus, we quantified acute cerebral microbleeds in hemibrains via H&E staining. Consistent with previous studies (Yakushiji et al., 2020, Hur et al., 2018, Han et al., 2015), we observed a significant increase in the number of microhemorrhages in Tg2576 mice (**Figure 3E, F**). In accordance with restoration of BBB breakdown by mTOR attenuation (**Figure 3A, C**), the number of microhemorrhages was significantly reduced in rapamycin treated Tg2576 mice to the point that it was not significantly different than WT + vehicle treated animals (**Figure 3E, F**). Thus, we demonstrate that mTOR drives BBB breakdown and acute cerebral microbleeds in aged Tg2576 mice with high fibrillar Aβ load.

### mTOR attenuation regulates proteins involved in the maintenance of the BBB

Prior work demonstrates tight junction protein modification underlies BBB breakdown (Biron et al., 2011, Van Skike et al., 2018). To assess changes in tight junction proteins, we examined the expression of claudin-5 and zonula occludens-1 (ZO-1), well-established tight junction proteins, by immunofluorescent confocal microscopy and measured immunofluorescent intensity. Claudin-5 (**Figure 4A-B**) and ZO-1 (**Figure 4C-D)** are upregulated in rapamycin-treated Tg2576 mice compared to Tg2576 vehicle-treated mice (p<0.0001 and p=0.04, respectively). The upregulation of claudin-5 and ZO-1 likely contribute to the improved BBB function we observed Tg2576 mice treated with rapamycin. Finally, MMP9 expression was assessed since increased expression is associated BBB breakdown (Weekman et al., 2016, Spampinato et al., 2017). MMP-9 levels were increased in vascular segments with fibrillar Aβ in Tg2576 mice relative to wildtype vehicle-treated mice (p=0.0306) (**Figure 4E-F**). This increase in MMP9 was rescued by mTOR attenuation with rapamycin in AB+ vessels (p=0.0138). This shows that mTOR attenuation rescues levels of tight junction proteins and an extracellular matrix degrading protein, possibly explaining our finding of rescued BBB function.

**Figure 4.**
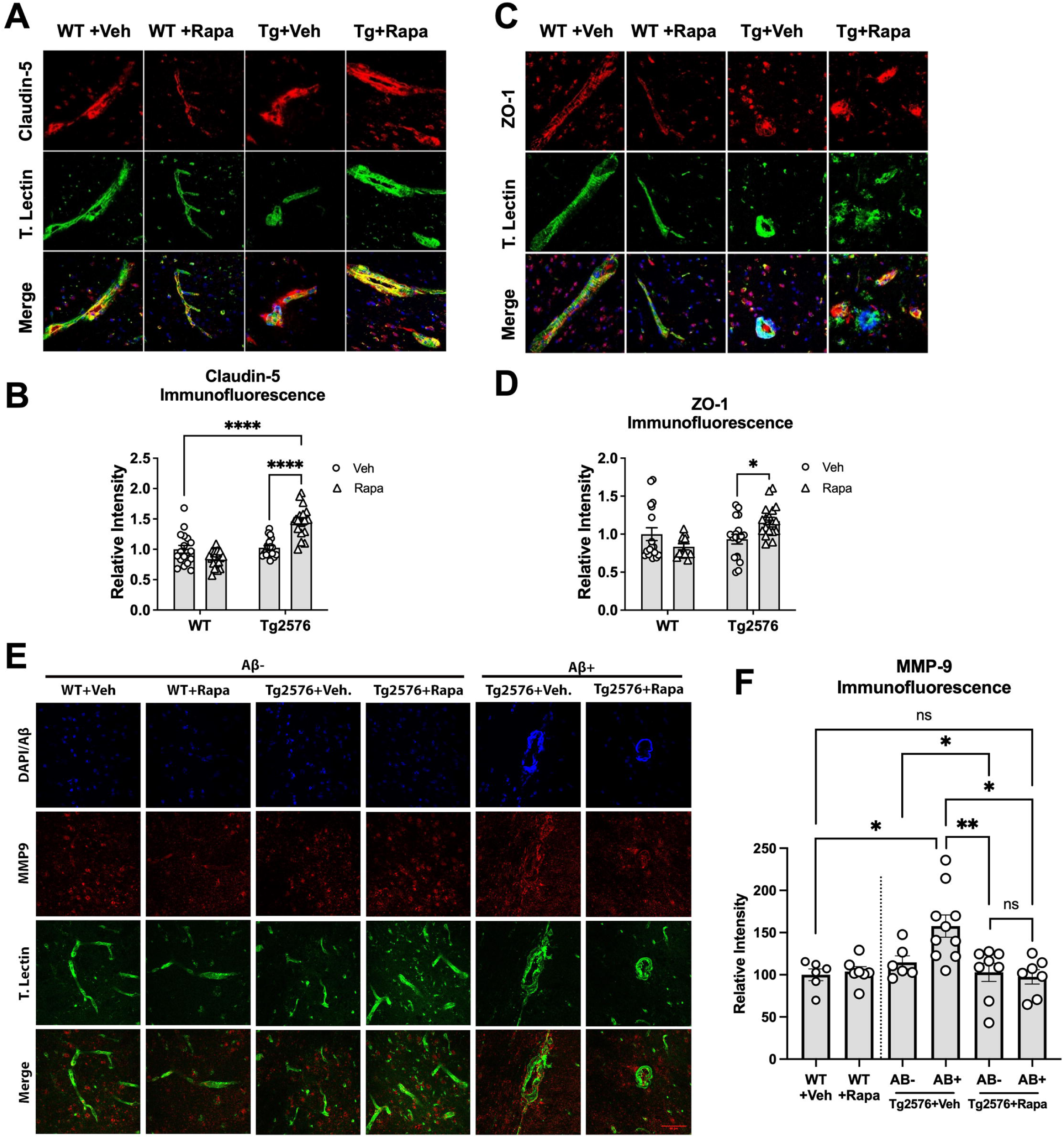
Attenuation of mTOR upregulates tight junction protein expression and downregulates MMP-9 expression. **(A)** Representative immunofluorescent images of claudin-5 (red) and brain vasculature (tomato-lectin, green). **(B)** mTOR inhibition in Tg2576 mice upregulates immunofluorescent intensity of claudin-5 (**** Tukey’s post hoc q(62)>8.83, p<0.0001 versus WT+Vehicle and Tg2576+Rapa groups). **(C)** Representative immunofluorescent images of ZO-1 (red) and brain vasculature (tomato-lectin, green). **(D)** Immunofluorescent intensity of ZO-1 is increased in Tg2576 mice by rapamycin treatment (*q(62)=3.82, p=0.04). (K) Chronic mTOR attenuation in Tg2576 mice produces a marginal increase in microvascular expression of ZO-1 (q(14)=3.71, p=0.08). **(E)** Representative immunofluorescent images of MMP-9 (red), brain vasculature (tomato-lectin, green), and nuclei/ Aβ (DAPI/Methoxy-X04). (F) Rapamycin reduced immunofluorescent intensity of MMP-9 in Aβ+ and Aβ-regions of Tg2576 mice relative to Aβ+ and Aβ-regions vehicle-treated controls, respectively.

### Chronic mTOR attenuation negates NVC deficits by restoring the nNOS-dependent component of the NVC response

Normal brain function is dependent on cerebral blood flow (CBF) to deliver a continuous supply of oxygen and glucose, which must rapidly increase during neuronal activity to meet the energy requirements of active neurons (Claassen et al., 2021). Regional increases in CBF are linked to neuronal activity through a mechanism known as neurovascular coupling (NVC). Changes in NVC can significantly impact cognitive function (Iadecola, 2013, Sorond et al., 2013, Tarantini et al., 2015), and NVC deficits in patients with AD(Hock et al., 1997, Rombouts et al., 2000, Rombouts et al., 2005, Girouard and Iadecola, 2006) are recapitulated in various mouse models of the disease (Shin et al., 2007, Lourenço et al., 2017, Park et al., 2014, Tarantini et al., 2017b, Van Skike et al., 2021).

Neuronal nitric oxide synthase (nNOS) has a prominent role in the regulation of NVC through the production of NO, released by neurons in an activity-dependent manner(Dirnagl et al., 1993, Cholet et al., 1996, Buerk et al., 2003). In addition, increasing evidence supports that NO derived from endothelial nitric oxide synthase (eNOS) also contributes to NVC through retrograde propagation of vasodilation(Toth et al., 2015a, Chow et al., 2020, Chen et al., 2014). We have previously demonstrated that unique contributions of nNOS and eNOS to neurovascular coupling can be independently isolated through the serial application of NOS inhibitors (Van Skike et al., 2021). Therefore, to define a potential role of reduced nNOS and eNOS activity in NVC deficits of Tg2576 mice, we measured evoked CBF within the barrel cortex in response to whisker pad stimulation as previously published (Van Skike et al., 2021).

In agreement with previous studies (Tarantini et al., 2017a), CBF increases in the somatosensory barrel cortex in response to whisker pad stimulation were reduced in 12-month-old Tg2576 mice (**Figure 5A, D**). Impaired NVC, however, was negated by mTOR attenuation in rapamycin-treated Tg2576 mice (**Figure 5A, D**). Neuronal NOS-derived nitric oxide contributes significantly to NVC responses (Cholet et al., 1996, Dirnagl et al., 1993) and we have previously demonstrated that mTOR inhibits nNOS to produce NVC impairments in the hAPP(J20) model of AD (Van Skike et al., 2021). Thus, to define the potential role of damage to specific NVC mediators in AD-like NVC impairment and determine the impact of mTOR attenuation of NO-dependent contributions to NVC and how each may drive NVC in Tg2576 mice, we measured CBF changes in the exposed barrel cortex after continuous superfusion (Dirnagl et al., 1994, Zhang et al., 1995, Leithner et al., 2010, Mishra et al., 2016, Toth et al., 2015b) of artificial cerebrospinal fluid (aCSF) first, to measure baseline (maximal) NVC responses; then after continuous superfusion of NOS inhibitors with different isoform specificity [i.e. nNOS (Beamer et al., 2012, Sousa et al., 2015, Bonvento et al., 2000, Burke and Buhrle, 2006, Stefanovic et al., 2007, Yang et al., 1999, Van Skike et al., 2021) versus all NOS (Leithner et al., 2010, Iadecola et al., 1993, Dirnagl et al., 1994, Van Skike et al., 2021)], to define specific contributions of NOS to NVC. The contribution of NOS activity to NVC responses was measured as the magnitude of the decrement in evoked CBF observed in the presence of each inhibitor [i.e. the magnitude of the L-NPA-sensitive response provided a measure of the contribution of nNOS; the magnitude of L-NPA + L-NAME-sensitive response, measured subsequently, provided a measure of the remaining, non-nNOS, eNOS contribution].

**Figure 5.**
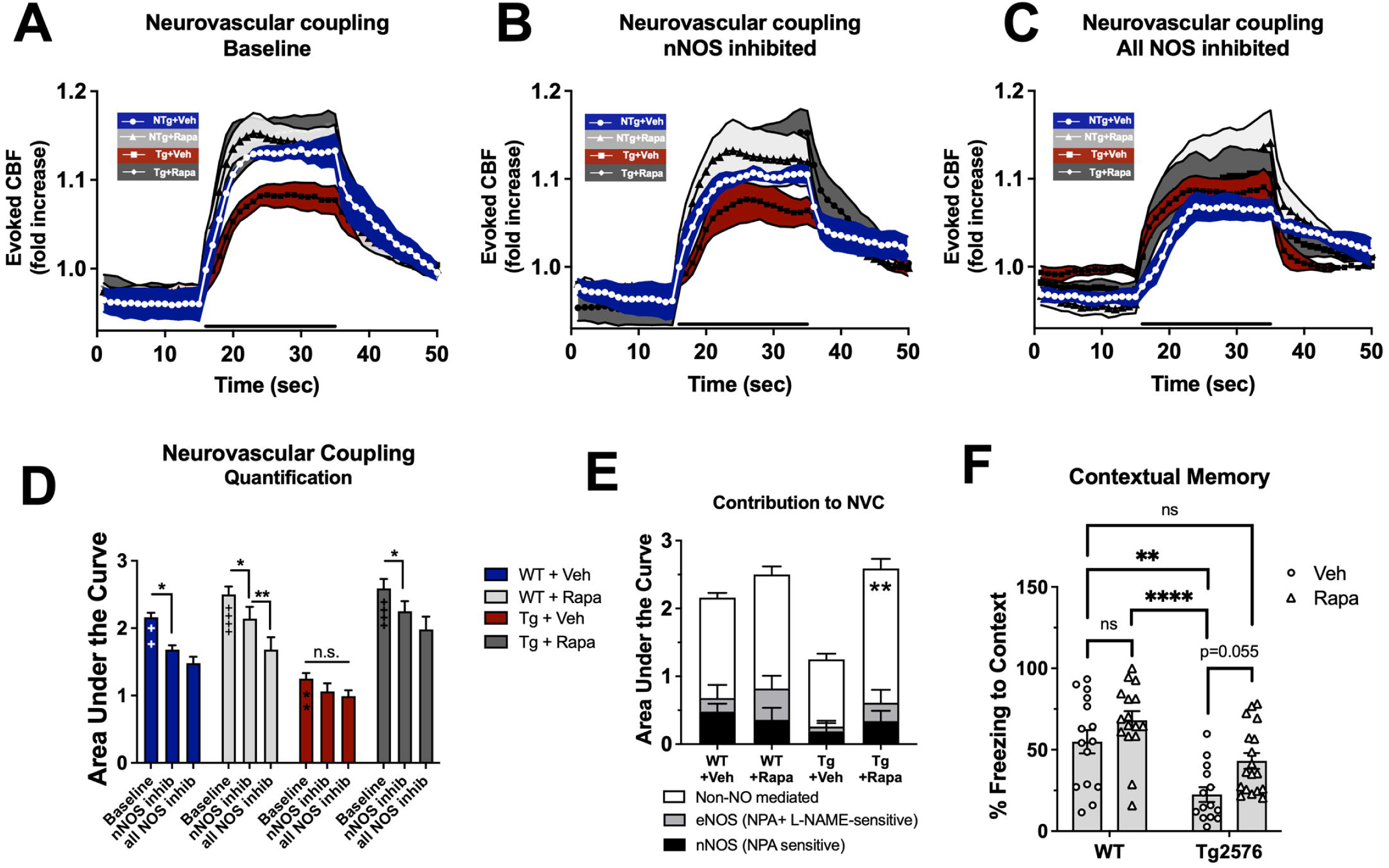
Inhibition of mTOR improves neurovascular coupling and contextual memory in Tg2576 mice. Long term attenuation of mTOR reverses neurovascular (NVC) impairments in 12-month-old Tg2567 mice. (**A–C**) Fold change in cerebral blood flow (CBF) relative to baseline are the mean ± SEM of *n* = 4 mice per group during whisker pad stimulation (30 s, bold black line). Stimulations were conducted sequentially in the presence of **(A)** aCSF [vehicle (Veh); **(B)** L-NPA to inhibit nNOS specifically, and **(C)** L-NPA+L-NAME to inhibit all NOS. The difference in CBF response using different these inhibitors allows for the measurement of the contribution of the nNOS [CSF – L-NPA] and eNOS [L-NPA – (L-NPA+L-NAME)], and the NOS-independent component of NVC [L-NPA+L-NAME]. (**D**) NVC responses during 30s whisker stimulations calculated as area under the curve defined as an increase relative to baseline (i.e., only upward peaks). Baseline NVC impairments in 12-month-old Tg2576 mice relative to WT (*q*_(12)_ = 8.501,^++++^*p*= 0.0003) are negated by rapamycin treatment (Rapa; *q*_(12)_ = 12.52, \*\*\*\**p* < 0.0001). Inhibition of nNOS with L-NPA significantly reduces NVC in WT mice (*q*_(36)_ = 2.580, \**p* = 0.014), but not in vehicle-treated Tg2576 mice (*q_(36)_ =* 4.407*, p=* 0.3140). These data suggest that Tg2576 mice are deficit in the nNOS component of NVC that is restored by rapamycin treatment, indicated by a significant inhibition of NVC in the presence of L-NPA (*q*_(36)_ = 1.935 vs baseline, \**p* < 0.0001), and in the presence of L-NPA+L-NAME (inhibiting all remaining NOS activity; i.e., eNOS) (*q*_(36)_ = 1.935 vs baseline, \**p* < 0.0001). The remaining NVC responses in the presence of L-NAME was not significantly different versus L-NPA alone in WT (*q*_(36)_ =0.000, *p* = 0 >0.9999), or rapamycin-treated Tg2576 mice (L-NPA+L-NAME vs L-NPA, *q*_(36)_ = 1.451, \**p* = 0.1554). **(E)** Contributions of nNOS, eNOS, and NOS-independent components to total NVC. There was an overall decrease in the NVC responses in Tg2576 mice, meanwhile deficits in NOS-independent NVC in Tg 2576 mice are negated by mTOR inhibition in rapamycin-treatment (*q*_(36)_ = 6.932, \*\**p* =0.0011). (**F)** Contextual fear memory deficits in Tg2576 mice (***q(65)=5.87, p=0.0006 versus WT+Veh) are significantly improved with mTOR attenuation via rapamycin (*q(65)=3.67, p=0.05), and are negated relative to WT (q(65)=2.42, p=0.33 versus WT+Veh). Tukey’s multiple-comparisons tests. Unless otherwise indicated, asterisks (*) in the figure represent a significant difference relative to Tg2576+Veh and pluses (+) represent a significant difference relative to WT+Veh.

Thus, to define a potential role of reduced nNOS activity in NVC deficits of Tg2576 mice, we used a selective inhibitor of nNOS, L-NPA at 200 nM, a concentration that effectively inhibits nNOS but does not impact eNOS-dependent responses *in vivo* (Van Skike et al., 2021). We found that superfusion of pial vasculature with 200 nM L-NPA significantly blunted CBF increases from somatosensory stimulation in WT animals **(Figure 5B, D),** however, NVC in Tg2576+Veh was insensitive to nNOS inhibition, indicating a significant impairment in nNOS activity in Tg2576 mice that was negated with mTOR inhibition (**Figure 5B, D**). Subsequent superfusion of L-NPA together with a general inhibitor of NOS (L-NAME, **Figure 5C**) only significantly blunted the CBF increase in WT mice treated with rapamycin, perhaps suggestive of enhanced eNOS activity.

To compare the magnitude of NOS contribution to the NVC response among the different groups of mice, using the area under the curve for each of the NVC responses **(Figure 5D**), we calculated the relative contribution nNOS-and eNOS-dependent NVC mechanisms (**Figure 5E**). Although inhibition of nNOS significantly reduced the NVC response relative to baseline (**Figure 5D**), the independent contribution of nNOS and eNOS to overall neurovascular coupling is not significant (**Figure 5E**), suggesting that non-NO mediated components of NVC underlie the impaired NVC response in Tg2576 mice. Unlike in hAPP(J20) mice, where mTOR drives nNOS and eNOS dysfunction and overall NVC deficits (Van Skike et al., 2021), the restoration of NVC by rapamycin in the Tg2576 model of AD suggests that mTOR contributes mainly to non-NO mediated components of NVC deficits (**Figure 5E**).

### mTOR drives hippocampal-dependent contextual memory impairment in Tg2576 mice

Cerebrovascular function is significantly linked to cognitive function (REF). Therefore, to determine whether the inhibition of mTOR with rapamycin was able to restore cognitive function in Tg2576 mice, we measured hippocampal-dependent contextual memory using the fear conditioning paradigm. Consistent with prior studies (Rodriguez-Rivera et al., 2011, Dineley et al., 2002, Comery et al., 2005, Taglialatela et al., 2009, Nobili et al., 2017), 10-month-old Tg2576 mice showed significant contextual memory impairment relative to WT littermates (**Figure 5F**). Attenuation of mTOR, however, negated contextual memory impairment in rapamycin-treated Tg2576 animals to a level indistinguishable from that of WT littermates (**Figure 5F**). Additionally, mTOR attenuation in WT animals did not alter contextual memory (**Figure 5F**). These data indicate that contextual memory impairments in Tg2576 mice are driven by mTOR, likely through the mechanisms we identified in this study including decreased vascular Aβ burden, improved vascular reactivity, BBB maintenance, and improved NVC.

## Discussion

Cerebrovascular dysfunction, a central driver of AD pathogenesis (Kisler et al., 2017, Girouard and Iadecola, 2006, Lecrux and Hamel, 2011, Zlokovic, 2011, Iturria-Medina et al., 2016), is recapitulated in various mouse models of AD amyloidopathy (Winkler et al., 2015, Merlini et al., 2011, Klakotskaia et al., 2018, Jullienne et al., 2023), including hAPP(J20) mice (Tong et al., 2009, Nicolakakis et al., 2008, Ongali et al., 2014, Royea et al., 2017, Royea et al., 2020, Van Skike et al., 2018, Lin et al., 2013), and in models of tauopathy (Bennett et al., 2018, de Jong and Jepps, 2018, Hussong et al., 2023) and multiple/mixed AD pathologies (Joo et al., 2017, Lourenco et al., 2017). In the present study, we choose the Tg2576 mouse model to investigate the mechanisms by which mTOR regulates CAA-related vascular dysfunction because of its accumulation of Aβ fibrils on the cerebrovasculature.

We find that mTOR drives the deposition of fibrillar Aβ on the cerebral vasculature (**Figure 2A-C)**. However, mTOR does not drive the accumulation of Aβ in the brain parenchyma. Attenuation of mTOR with rapamycin led to a significant decrease in vascular Aβ fibrils, without changing vascular density in the measured fields. These data suggest that the reduction is likely due to a specific decrease in vascular Aβ accumulation without altering local vascular density. It should be noted that the small imaging window utilized in the current study does not preclude an overall change in vascular density from the whole brain, as has previously been reported in the J20 mouse model of AD (Lin et al., 2013), however, there was no change in vascular density within the imaging region. In fact, the aged Tg2576 mice have been reported to have increased microvascular density, due to increased angiogenesis and hypervascularization (Biron et al., 2011).

Additionally, the enhancement in endothelium-dependent vascular reactivity as a result of mTOR attenuation was observed in all genotypes and both in Aβ+ and Aβ-vessels, suggesting that the restoration of cerebrovascular reactivity by mTOR attenuation is not simply a consequence of decreased Aβ fibrils accumulation in cerebrovasculature, but modulation of mTOR. We have previously shown that mTOR attenuation with rapamycin improves peripheral blood flow in response to vasodilators in a mouse model of AD, indicating potential beneficial effects of mTOR attenuation on endothelial function and vascular reactivity (Van Skike et al., 2023).

Moreover, our data suggest that the mechanisms by which mTOR attenuation restores cerebrovascular reactivity in Aβ+ vessel segments are downstream of Aβ-induced cerebrovascular dysfunction. One candidate for this dysfunction is deficient eNOS signaling. Although, activation of eNOS in the microvasculature could not be assessed in these animals, we have previously shown that eNOS activation in the microvasculature peaks 1-3 minutes after ACh administration and subsequently returns to baseline, prior to maximal vasodilation of the endothelium (Lin et al., 2013), thus, the timeline of our present studies extends beyond the time period required for accurate assessment of ACh-induced eNOS phosphorylation.

We identified a significant positive correlation between vascular Aβ burden and BBB breakdown in Tg2576 mice (r=0.80, p=0.003, **Figure 3D**). BBB permeability and Aβ deposition have been shown to interact in the Tg2576 model, with some reports implicating BBB permeability as causal to Aβ deposition (Ujiie et al., 2003), and others finding that Aβ and CAA cause BBB disruption in this model (Biron et al., 2011). Through mTOR attenuation, we find a reduction in vascular Aβ lesions (**Figure 2**) and improvement in BBB integrity (**Figure 3C**). Since rapamycin affected both measures, a causal relationship cannot be determined, however, these factors likely interact in a negative feedback loop to contribute to overall disease burden. We also observed that the cerebral microbleeds that occur in Tg2576 mice are rescued by rapamycin, providing additional evidence of improved BBB function with mTOR attenuation. Furthermore, we observed that expression of tight junction proteins claudin-5 and ZO-1 increased with rapamycin treatment (**Figure 4A-D**). Levels of the MMP9 decreased in Aβ+ vessels, suggesting that mTOR regulates cerebrovascular tight junction protein modification in the Tg2576 model of AD with high vascular Aβ load, potentially underlying the observed BBB breakdown and cerebral microbleeds (**Figure 4E-F**).

We found that NVC responses are reduced in Tg2576 mice but are rescued by rapamycin **(Figure 5A**). Our data show that the non-NO mediated portion of NVC responses are negatively affected by mTOR signaling in Tg2576 mice. We saw that Tg2576 mice were insensitive to nNOS (L-NPA inhibited response) and eNOS (L-NAME inhibited response) inhibition, which were improved by mTOR attenuation. While L-NAME does not specifically inhibit eNOS, but inhibits all NOS, a contribution of L-NAME-sensitive iNOS activity to NVC is unlikely because no role for iNOS in NVC or mechanisms of regulation for this NOS isoform that are responsive to changes in neuronal activity have been documented. Thus, the magnitude of the non-nNOS, L-NAME sensitive NVC response in our studies provided a measure of eNOS activity. The magnitude of the remaining NVC response after inhibition of all NOS provided a measure of non-NO mediated contributions to NVC. Thus, we conclude that mTOR contributes to non-NO mediated NVC defects in the Tg2576 model of AD with CAA.

We (Spilman et al., 2010, Lin et al., 2013) and others (Caccamo et al., 2014) have shown that mTOR attenuation prevents (Caccamo et al., 2014, Spilman et al., 2010, Majumder et al., 2011) and reverses (Lin et al., 2013, Van Skike et al., 2021) cognitive deficits in the hAPP(J20), Tg2576, 3xTg-AD mouse models of AD. To determine whether mTOR contributes to cognitive impairment in Tg2576 mice and to determine the potential involvement of mTOR-driven cerebrovascular dysfunction to AD-like deficits, we performed contextual fear conditioning. This cognitive test was chosen because the hyperactivity and increased locomotion of the Tg2576 model (Lalonde et al., 2012, Gil-Bea et al., 2007) makes it difficult to perform other cognitive tasks. We found that contextual memory impairments are also negated by rapamycin, indicating a broad beneficial effect of rapamycin across many different models of of AD, VCI, and normative aging(Van Skike et al., 2021, Jahrling et al., 2018, Van Skike et al., 2018, Van Skike et al., 2020, Caccamo et al., 2014, Lin et al., 2013, Majumder et al., 2011, Spilman et al., 2010).

We find that mTOR contributes to cerebrovascular dysfunction, including decreased vascular reactivity, BBB breakdown, and impaired neurovascular coupling, by driving the accumulation of fibrillar Aβ on the cerebral vasculature. Attenuation of mTOR by rapamycin improved these parameters, as well as contextual memory. This supports the therapeutic potential of using mTOR inhibitors to treat CAA and AD.

## Author Contributions

Conceptualization: CEVS, SAH, KTD, VG. Investigation: CEVS, SFH, SAH, ND, JBJ. Formal analysis: CEVS, SFH, LRM, SAH, HM, ACM, ND, JBJ. Writing-original draft: CEVS. Writing-editing and reviews: LRM, HM, ACM, SAH, VG. Funding acquisition: KTD and VG.

## Supporting information

Publication License from BioRender

Publication License from BioRender

## Acknowledgements

We acknowledge the following funding support: NIH National Institute on Aging (NIA) 1R01AG057964-01 (Galvan) and US Department of Veterans Affairs I01 BX002211-01A2 (Galvan), the Oklahoma Nathan Shock Center on Aging (Galvan) NIA 2- P30AG050911-11, San Antonio Medical Foundation (Galvan), the JMR Barker Foundation (Galvan), and a William & Ella Owens Medical Research Foundation Grant. Dr. Galvan is the recipient of a Research Career Scientist award, IK6 BX006026, from the US Department of Veterans Affairs. Dr. Galvan acknowledges the generous support from the Robert L. Bailey and daughter Lisa K. Bailey Alzheimer’s Fund in memory of Jo Nell Bailey. We also recognize NIA Training Grant T32AG021890 and the Alzheimer’s Association AARF-17-504221 (Van Skike), US Department of Veterans Affairs Biomedical Laboratory Research and Development Service IK2 BX003798-01A1 and a pilot award from the Oklahoma City Veterans Health Care System (Hussong). K.T.D. was supported by NIH/NINDS R01NS094557. These studies used the services of the Healthspan and Functional Assessment Core and the Pathology Core of the San Antonio Nathan Shock Center of Excellence in the Biology of Aging (NIH/NIA 2 P30 AG013319-21). Two-photon images were generated in the Core Optical Imaging Facility, which is supported by UTHSCSA, NIH-NCI P30 CA54174 (CTRC at UTHSCSA) and NIH-NIA P01A; San Antonio Nathan Shock Center Pathology Core.

